# Characterization of LXR-activating nanoparticle formulations in primary mouse macrophages

**DOI:** 10.1101/746784

**Authors:** Tyler KT Smith, Zaina Kahiel, Nicholas D LeBlond, Peyman Ghorbani, Eliya Farah, Marceline Cote, Suresh Gadde, Morgan D Fullerton

## Abstract

Activation of the transcription factor liver X receptor (LXR), has shown to be efficient at curbing aberrant lipid metabolism and inflammation. While small molecule delivery via nanomedicine has promising applications for a number of chronic diseases, there remain questions as to how nanoparticle formulation might be tailored to suit different tissue microenvironments and aid in drug delivery. In the current study, we compared the drug delivery capability of three nanoparticle (NP) formulations encapsulating the LXR activator, GW-3956. We observed little difference in the base characteristics of standard PLGA-PEG NP when compared to two redox-active polymeric NP formulations (DD and DB). Moreover, we also observed similar uptake of these NP into primary mouse macrophages. After an initial acute uptake period and using the transcript and protein expression of the cholesterol efflux protein ATP binding cassette A1 (ABCA1) as a readout, we determined that while the induction of transcript expression was similar between NPs, treatment with the redox-sensitive DB formulation resulted in a higher level of ABCA1 protein 24 h after the removal of the drug-containing NPs. Our results suggest that NP formulations responsive to cellular cues may be an effective tool for targeted and disease-specific drug release.

## 1. Introduction

Liver X receptors (LXR), a family of nuclear receptors, are attractive targets for therapeutic intervention in cardiovascular, metabolic and inflammatory diseases, owing to their regulation of several metabolic pathways including bile acid, carbohydrate, and lipid metabolism [1,2]. LXR activation protects against aspects of atherosclerosis via the induction of reverse cholesterol transport in macrophages. This stimulates cholesterol efflux by transcriptionally upregulating the efflux machinery, ATP-binding cassette (ABC)A1 and G1 [3]. Activation of LXR has also exhibited anti-inflammatory properties by trans-repressing NF-κB and enhancing macrophage efferocytosis. In addition to macrophages, activation of LXR has been shown to modulate inflammatory gene expression in many other cell types, such as T and B lymphocytes, microglia, astrocytes, and dendritic cells [4,5]. Induction of LXR signaling has been associated with decreased inflammation in mouse models of several acute and chronic diseases such as Alzheimer’s disease, lupus-like autoimmunity, experimental stroke, infection with Mycobacterium tuberculosis and atherosclerosis [6–8]. In this context, several synthetic LXR agonists have shown promise in the development of therapeutics for atherosclerosis and inflammatory diseases by promoting cholesterol efflux and inhibiting inflammation [9]. However, clinical translation of LXR-based therapeutic strategies have been dampened by hypertriglyceridemia and hepatic steatosis, which are caused by LXR-mediated induction of lipogenesis in the liver [10,11]. Therefore, innovative strategies that reduce the adverse side effects of LXR activation while maintaining efficacy are necessary for the development of LXR-based therapeutics [12–14].

Nanotechnology applications in medicine, nanomedicines, have shown the clinical impact on different disease therapies via drug delivery, imaging, and diagnostics [15]. In general, nanoparticle-based drug delivery platforms improve the pharmacokinetic profile of drugs, decrease toxicity, and deliver drugs in a tissue and organ specific manner [16].Advances in nano-biomaterials synthesis has enabled the development of nanoparticles (NP) that can encapsulate single and/or multiple drugs with different physicochemical properties and deliver them across multiple biological barriers in a targeted and controlled-release manner [17]. While there has been significant progress made toward use of nanomedicines in human cancers in recent years, nanotechnology applications are also well explored in cardiovascular, inflammation and infectious diseases. Spatiotemporal delivery of pro-inflammation resolution mediators such as Ac2-26 peptide, or anti-inflammatory cytokine IL-10 to atherosclerotic plaques using polymeric NPs improved the bioavailability of the agents and enhanced the therapeutic efficacy [18–20]. NP platforms were also used for modulating the polarity of monocytes and macrophages toward a less inflammatory phenotype and promoting the resolution of inflammation, ultimately preventing plaque destabilization and markers of rupture [12,21].

The majority of the NP platforms used in cardiovascular nanomedicines are based on polymeric nanocarriers. In general, a therapeutic payload from the polymeric NPs can be released via diffusion, erosion, and degradation [16]. Additionally, nanocarriers can be designed to release payloads via the response to either endogenous or exogenous stimuli. When compared to exogenous stimuli (magnetic field, radiation, ultrasound, etc.), endogenous stimuli prove to be more interesting, as they can be specific to disease-related pathological changes (pH differences between tumor microenvironment and normal cells, differential redox state, etc.) [22–24]. In the presence of intracellular reducing agents, various organic materials and molecules containing disulfide bonds can be bio-reduced to their thiols counterparts. Using this strategy, biomaterials with disulfide backbones were developed for reduction-responsive drug delivery nanocarriers and tested in tumor models [25,26]. However, although in the context of cancer biology this has been addressed, there have been no other studies that have assessed the effects of intracellular reducing agents on payload release and LXR activation in macrophages.

Therefore, in this study, we sought to perform a systematic evaluation of differential drug release effects on LXR activation in primary murine bone marrow-derived macrophages (BMDM). Effectively, we aimed to test whether basal levels of intracellular reducing agents were able to modulate redox-responsive NPs. Towards this, we developed three different NP platforms, traditional FDA approved polymeric NPs (Poly lactic acid-co-glycolic acid-polyethylene glycol, PLGA-PEG), NPs for slow and controlled release, as well as two redox-responsive disulfide backboned NPs named DB (dithiodibutyric acid backbone) and DD (dodecanedioic acid backbone). To activate LXR, we encapsulated the synthetic LXR agonist, GW-3965 (GW) into our NPs and tracked the transcript and protein levels of the LXR target gene, ABCA1.

## 2. Materials and Methods

### 2.1 Materials

PLGA-COOH is purchased from lactel polymers, NH_2_-PEG-COOH was purchased from Lysan Bio, Inc. PEG-COOH, HO-PEG-OH were purchased from BDH chemicals. 4,4’-dithiodibutyric acid (DBA, 95% purchased from Sigma-Aldrich), Dodecanedioic acid (DDA, >98% purchased from Fluka Chemika), 1,6-hexanediol (16HD, purchased from Sigma Aldrich), 2-dihydroxyethyl disulfide (2HDS, technical grade, purchased from Sigma Aldrich), anhydrous dichloromethane (DCM, 99.8%, purchased from Anachemia), methanol (MeOH, HPLC grade, 99.9%), N-hydroxysuccinimide (NHS, 98%), N-(3-dimethylaminopropyl)-N′-ethylcarbodiimide hydrochloride (EDC, commercial grade), N,N-diisopropylethylamine (DIEA, 99%), N,N-dimethylformamide (DMF, 99.9%), dimethylsulfoxide (99.5%), 4’(dimethylamino)pyrimidine, (DMAP, 99%, purchased from Sigma Aldrich), N,N’-dicyclohexycarbodiimide (DIC, 99%, purchased from Alfa Aesar), acetonitrile (can, >99.8%, purchased from Sigma Aldrich), and chloroform-D (CDCl3, purchased from Sigma Aldrich).

### 2.2 Polymers synthesis

PLGA-PEG polymer was synthesized as previously described [12].

### 2.3 Synthesis of DBA-16HD-PEG_2K_

1.00g of DBA (4.2 mmol) was dissolved in 5 mL of DMF and then placed in an ice bath. To this, 1.3 ml of DIC (2.2 equivalents) was added and then stirred for 30 min. Then, 0.610 g of 16HD (1.2 equivalents) was added following 1.0026 g addition of DMAP (2 equivalents). The solution was stirred overnight. Crude solutions were concentrated under vacuum and redissolved in DCM and washed with H_2_O to remove the byproducts of the reaction, dialyzed in DCM and proceeded to the next step. Next, 100 mg of PEG_2K_ (0.05 mmol) was added of the crude solution and placed in an ice bath under stirring conditions. In a slow, dropwise manner, 23 μl of DIC (0.15 mmol) was added to the solution and then it was stirred for 30 min. Then, 25 mg of DMAP (0.2 mmol) was added and then the solution was stirred overnight. The crude product was placed in a 3.5 K dialysis tube and dialyzed in 2:8 methanol: DCM solution and had the supernatant change every 6 h for a total of 24 h. The final product was characterized by GPC and ^1^H NMR spectroscopy. ^1^HNMR (CDCl3) □ in ppm: 4.01-4.08 (COO-C**H**_2_(CH_2_)_4_-C**H**_2_-OOC-), 3.64-3.59 (-OC**H**_2_C**H**_2_O-), 2.74-2.64 (-OOCCH_2_CH_2_C**H**_2_-SS-C**H**_2_CH_2_CH_2_COO-), 2.36-2.43 (-OOCC**H**_2_CH_2_CH_2_-SS-CH_2_CH_2_C**H**_2_COO-), 2.08-1.94 (-OOCCH_2_C**H**_2_CH_2_-SS-CH_2_C**H**_2_CH_2_COO-), 1.52-1.66 and 132-1.40 (COO-CH_2_(C**H**_2_)_4_-CH_2_-OOC-).

### 2.4 Synthesis of DDA-2HDS-PEG_4K_

500 mg of DDA (2.17 mmol) in DMF and kept in an ice bath and 670 μl of DIC (2.2 equivalents) was added dropwise and stirred for 30 min. 132 μl of 2HDS was then added to the solution and stirred for 3-4 h. To this reaction mixture 1.08 g of PEG_4K_ (0.27 mmol) was added, followed by 0.53 g of DMAP (4.33 mmol) added to the solution and stirred for 48 h. The crude product was dialyzed using 10 K MWCO dialysis membrane using 1:1 MeOH: DCM, for 12 h while the supernatant was replaced every 3 h. The final product was characterized by GPC and ^1^H NMR spectroscopy. ^1^HNMR (CDCl3) □ in ppm: 2HDS peaks: 4.34-4.22, 2.94-2.84; PEG: 3.72-3.54, DDA peaks: 2.43-2.38, 2.31-2.25, 1.65-1.53, 1.33-1.21.

### 2.5 GW-NPs synthesis and characterization

All GW-NPs were synthesized via a single step self-assembly process using the nanoprecipitation method. Briefly, polymers were dissolved in acetone or acetonitrile (10 mg/ml) and mixed with GW in acetonitrile in 10:1 w/w ratio and added to water dropwise. NPs were stirred at room temperature for 5-7 h and concentrated using 100 K MWCO pall centrifugal filters. NPs were washed twice with water and diluted 10-20 times in H2O or 1% PBS and characterized for their physicochemical properties. Hydrodynamic sizes and zeta potentials measurements were done using Zeta view and Malvern zetasizer. For TEM (transmission electron microscopy) imaging, freshly prepared NPs diluted 20 times and deposited 300 mesh carbon-coated copper grids. After grids dried, TEM imaging was done using FEI Tecnai G2 Spirit Twin electron microscope. The encapsulation efficiency of GW was estimated using HPLC, eluted with ACN: H2O gradient. [12] Encapsulation efficiencies were calculated as: [amount of GW in NPs/total amount of GW]*100. Stability of NPs were tested by incubating them for 1 hour in 5%, 10%, and 20% of fetal bovine serum (FBS). The NP size before and after incubation in serum was obtained with quasi-electric laser light scattering using a ZetaPALS dynamic light-scattering detector (15 mW laser, incident beam ¼ 676 nm; Brookhaven Instruments).

### 2.6 Mice

C57BL/6J mice were purchased from Jackson Laboratories (stock no. 000664) and bred in house. Mice were housed in a ventilated cage system and maintained on a 12 h light/12 h dark cycle with lights on at 0700 h with unlimited access to standard rodent chow (Envigo #2018) and water. All animal experimental procedures were in accordance with the guidelines and principles of the Canadian Council of Animal Care and were approved by the Animal Care Committee at the University of Ottawa.

### 2.7 Cell Culture

Bone marrow-derived macrophages were generated as previously described [27]. Briefly, mice were euthanized, tibias and femurs isolated, and the ends of each bone cut off. The tibia and femur from each leg were placed into a sterile 0.5 ml microfuge tube that had a hole punctured in the end with an 18-gauge needle, which was then placed inside of a 1.5 ml microfuge tube before the addition of 100 µl of DMEM (Wisent) to the 0.5 ml tube. Bone marrow cells were collected by centrifuging at 4,000 rpm for 5 min, resuspended, filtered, and plated in 80 ml of DMEM supplemented with 10% FBS (Wisent) and 1% penicillin/streptomycin (Thermo Fisher) in a T175 flask, and incubated at 37ºC in a humidified atmosphere at 5% CO2. After 4hr, cells were plated in 15 cm tissue culture dishes in the presence of 20% L929 medium (as a source of macrophage colony stimulating factor) and left to differentiate for 7-8 days. One day prior to the experiment, cells were lifted into suspension in the existing L929-supplemented DMEM by gently scraping and seeded into the appropriate plate for subsequent experiments.

### 2.8 GW Treatment

Cells were seeded at a density of 1.2 × 10^6^ cells/well in 6 well plates and 0.6 × 10^6^ cells/well in 12 well plates in DMEM supplemented with 10% FBS and 1% penicillin/streptomycin. After adherence, BMDM were treated with 5 µM GW in either free-drug form or encapsulated in NPs for a period of 90 min. Cells were then washed twice with PBS (Wisent) to remove drug and nanoparticle treatments, and complete DMEM was reapplied for the duration of the experiment, with the exception of the chronic GW dose, in which 5 μM GW was replenished for the entirety of the treatment period. All time points indicate time post 90-min treatment.

### 2.9 Western Blotting

Cellular lysates were prepared, and Western blotting and quantification were performed as previously described [27]. Anti-ABCA1 (1:1000; Novus) and anti-beta-actin (1:1000; Cell Signaling) primary antibodies were used with an HRP-conjugated anti-rabbit IgG secondary antibody (1:10000; Cell Signaling).

### 2.10 RNA Isolation, cDNA synthesis, and quantitative PCR

RNA was isolated using TriPure (Roche) according to the manufacturer’s instructions. Total RNA was DNase-I treated (ABM) and first strand synthesis was performed using OneScript Plus reverse transcriptase. cDNA was diluted 1:20 into ultrapure water, and mRNA expression of β actin and Abca1 was determined using TaqMan gene expression assays (Thermo Fisher Scientific). Relative expression was calculated using the ^ΔΔ^Ct method, as previously described [28].

### 2.11 Flow Cytometry

BMDM were treated with 7.5 x10^9^ NPs/well of Cy5.5-conjugated (Ex/Em: 675/694) NPs for 1.5 h in 6 well plates then washed twice with PBS. 5mM EDTA (Wisent) was added for 5 min at 37 °C to lift cells off the plate, after which they were transferred to a 96 well V-bottom plate (Life Science). BMDM were centrifuged at 360 x g for 7 min then resuspended in PBAE buffer [1% BSA (Thermo Fisher), 0.01% sodium azide, 1% EDTA) with DAPI (Invitrogen, 1:1000) in PBS]. Cells were then sorted using the BD LSRFortessa™ cell analyzer.

### 2.12 Microscopy

BMDM were treated with Cy5.5-conjugated NPs for 1.5 h then washed twice with PBS. 1% paraformaldehyde in PBS was added to the cells for fixation for 15 min at room temperature. The cells were then washed and simultaneously blocked and permeabilized with a 1% BSA and 0.1% Triton X-100 PBS solution for 30 min at room temperature. Cell were then stained with a LAMP1-eF450 (1:100; Ex/Em: 405 nm/450 nm; eBioscience) primary antibody in PBS for 30 min. BMDM were washed again and immediately prior to mounting were stained with 0.5 ml/million cells of PI-RNAse solution (Ex/Em: 493 nm/636 nm; BD). BMDM were imaged on the Zeiss LSM800 AxioObserver Z1. Images were later analyzed using ImageJ.

### 2.13 Statistical analyses

Unless otherwise stated, data represent mean ± SEM. Statistical analysis was performed using GraphPad Prism (v7.0) where a one-way ANOVA was used to determined significant differences. A Tukey posthoc test was used to determine significant differences (P<0.05) between treatments.

## 3. Results

### 3.1. Synthesis and characterization of polymers and nanoparticles

To compare the effects of GW release on LXR activation, we developed three different NPs (Figure. 1) with differential drug release properties. To this end, we synthesized polyester polymers with and without disulfide backbones. Di-block polymer PLGA-PEG was synthesized according to published procedures [12]. Briefly, the terminal carboxyl group of PLGA polymer was reacted with the amine moiety of PEG via a N-(3-dimethylaminopropyl)-N′-ethylcarbodiimide /N-Hydroxysuccinimide (EDC/NHS) coupling reaction. Polyester polymers named DB16HD and DDHP4, with disulfide bonds in their backbone were synthesized via a polycondensation reaction between diacids and diol groups, and capped with the PEG moieties. For this, 4,4’-dithiodibutyric acid was reacted with a slight excess of 1,6-hexanediol, using a N,N′-diisopropylcarbodiimide/4-dimethylaminopyridine (DIC/DMAP) condensation reaction. The crude product was purified and reacted with carboxylic acid groups on PEG to obtain the final product DB16HD (SI Figure 1).

**Figure 1:**
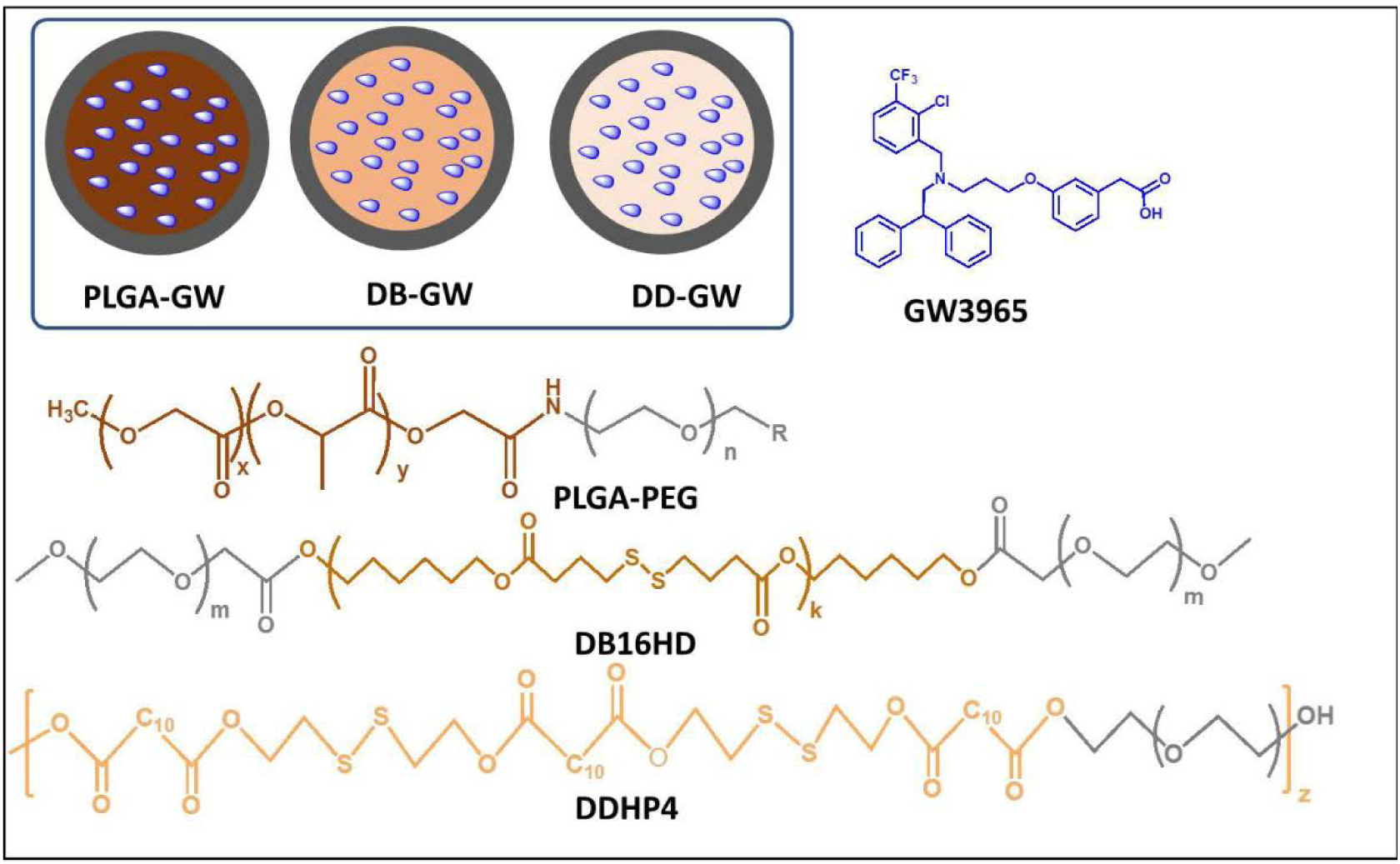
Schematic diagram of nanoparticles used in this study (inset), and chemical structures of polymers and GW-3965.

In the case of DDHP4, we employed a similar strategy, but used 2-dihydroxyethyldisulfide and dodecanedioic acid as a building blocks. All polymers were characterized by proton nuclear magnetic resonance spectroscopy (^1^H-NMR) and gel permeation chromatography (GPC). Next, we developed three different NPs containing GW via a single step self-assembly process using the nanoprecipitation method [12,29], to produce slow releasing PLGA-GW, as well as redox-reactive DB-GW (synthesized using DB16HD polymer) and DD-GW (synthesized using DDHP4 polymer) NPs. The appropriate ratios of polymers and GW were dissolved in water-miscible organic solvent acetonitrile and introduced to deionized water under constant stirring, forming a solid polymeric core containing GW and a PEG shell.

Upon synthesis, NPs were purified and characterized for their size, surface charge, and GW encapsulation efficiency. The hydrodynamic size measurement of NPs by both Dynamic Light Scattering (DLS) and ZetaView particles tracking indicated that the size was approximately 100-200 nm for all particles with narrow size distributions (SI Figure 2 and Figure 2A). All NPs were spherical in shape as shown by transmission electron microscopy imaging (Figure 2B) and had slightly less size due to the lack of the solvent hydration layer. All NPs also had a slightly negative surface charge with zeta potential values ranging from −3 to −7 mVs (Figure 2C). The encapsulation efficiencies of GW were determined by HPLC analysis of NPs, and were within the range of typical drug encapsulation, with 60-85% encapsulation efficiency [12–14]. The redox active NPs DB-GW and DD-GW encapsulated 67% and 85% of the original GW feed, and PLGA-GW NPs encapsulated 80% of GW in the NPs. We next explored the stability of NPs by incubating them in differing percentages of fetal bovine serum (FBS) at 37°C, to mimic physiological conditions. All NPs were stable within these mimicking conditions (Figure 2D).

**Figure 2:**
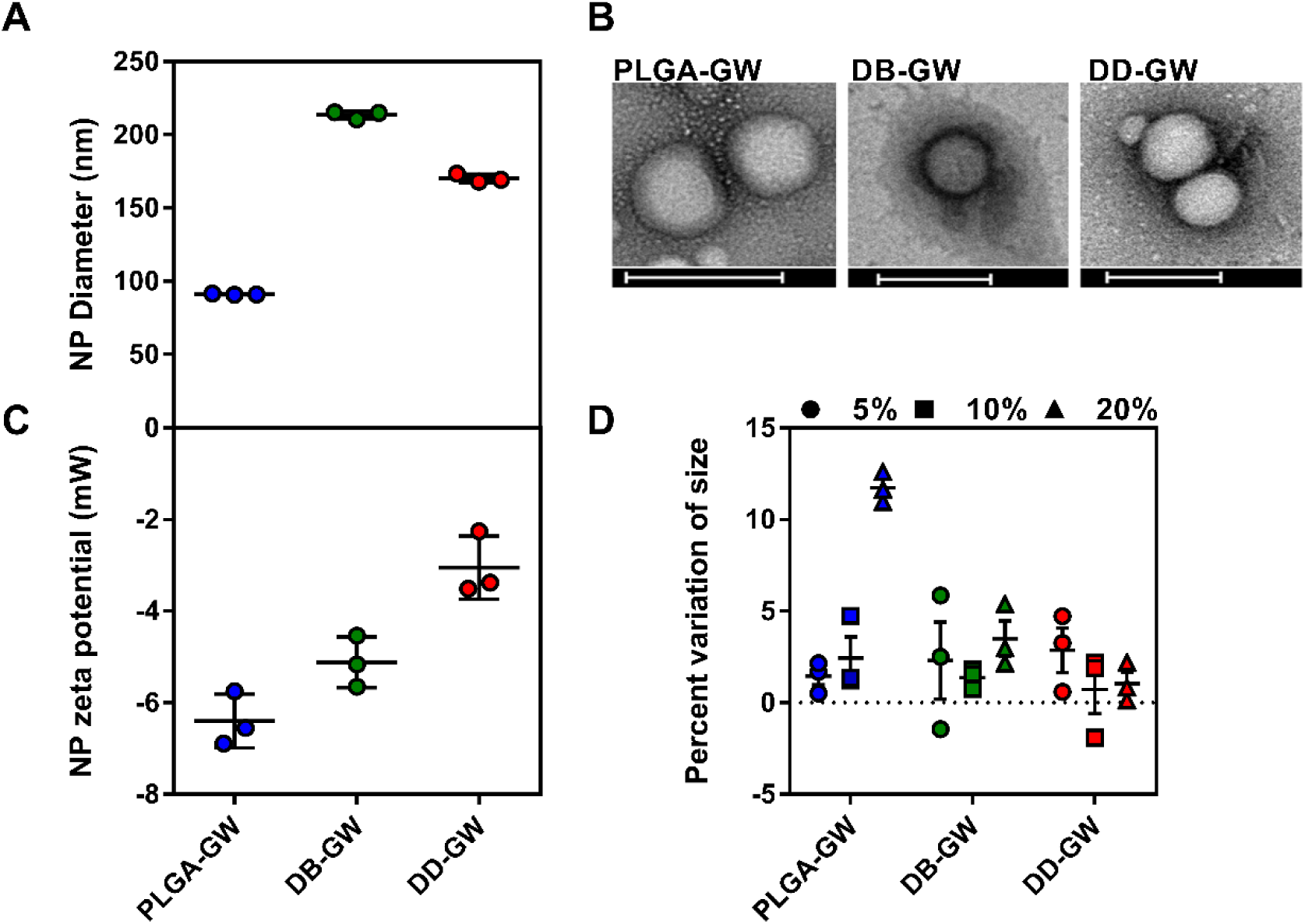
Nanoparticle characterization. A) Hydrodynamic size of three different GW-NPs measured by ZetaView and zetasizer (n=3). B) Transmission electron microscopy images of GW-NPs showing spherical structure, scale bar = 200 nm. C) Surface charge potentials of GW-NPs measured using zetasizer (n=3). D) Stability studies were performed by measuring the GW-NPs size, pre and 1 hour post incubation in 5%, 10%, and 20% FBS (n=3).

### 3.2 Cellular uptake experiments

In order to compare the effects of LXR activation by different NPs in a cellular system, we first studied if primary murine macrophages took up NPs at a similar rate. To this end, we developed NPs tagged with Cy5.5 dye by chemically conjugating Cy5.5 to the polymer. The Cy5.5-PLGA, Cy5.5-DB and Cy5.5-DD NPs were then analyzed by ZetaView to obtain NPs particle concentration in the solution. Cells were treated with Cy5.5-NPs in similar particle concentrations for 90 min, fixed and analyzed by flow cytometry to identify cells that had accumulated NPs. Our results indicated that the three NP formulations were taken up by macrophages at a similar rate (data not shown). Cells were also imaged by confocal microscopy with co-staining for LAMP1, a lysosomal marker. Consistent with the flow cytometry analyses, imaging determined that the uptake between the various NPs was similar (Figure 3).

**Figure 3.**
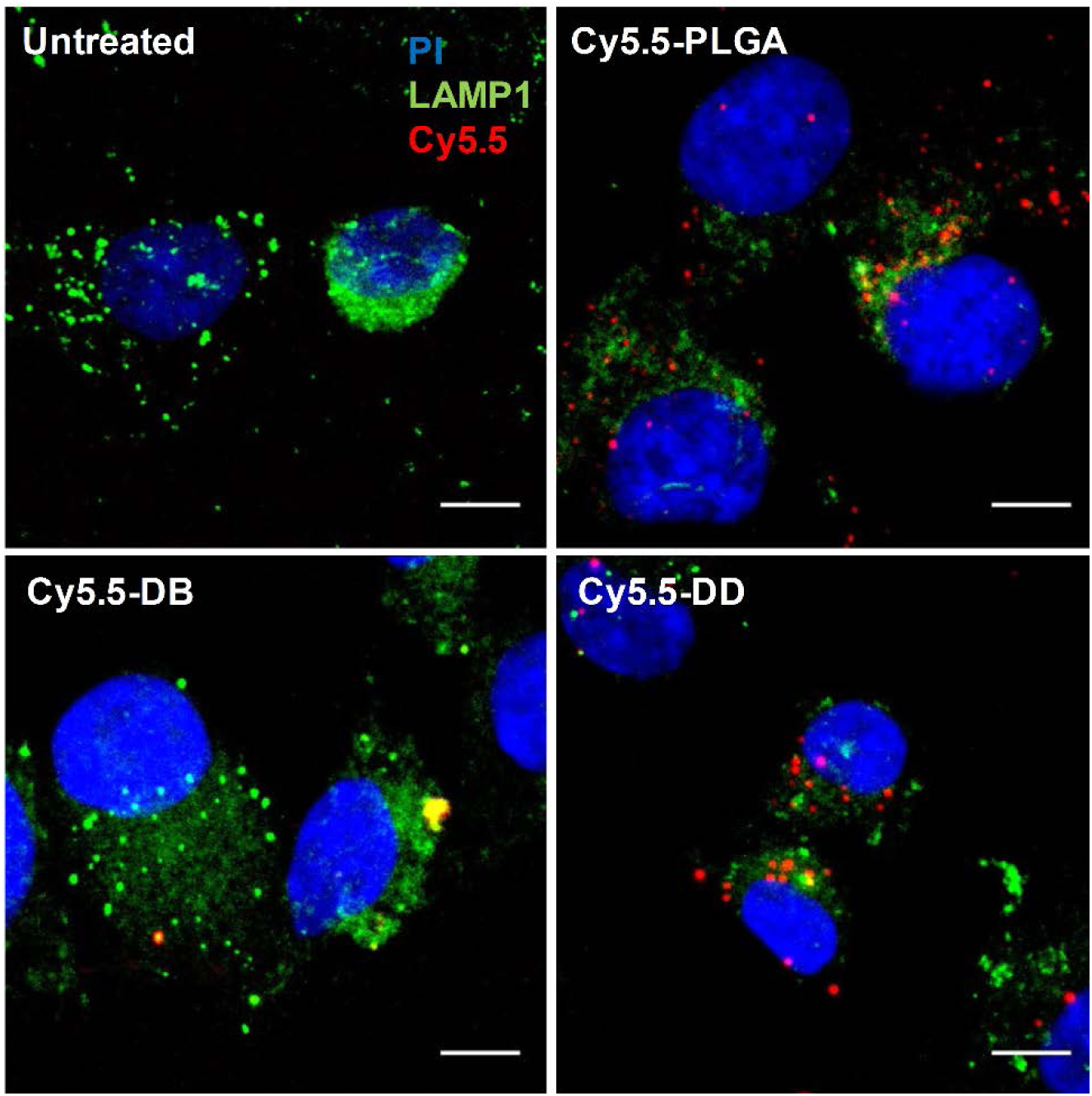
Nanoparticle formulations are effectively up-taken by macrophages and form punctate foci. WT BMDM were treated with 1.0×10^9^ Cy5.5-tagged nanoparticles (red) for 1.5 h, then fixed and stained with an anti-LAMP1 antibody as a lysosomal marker (green) and propidium iodide as a DNA stain (blue). Images representative of at least 5 fields of view from n=3. Scale bar represents 5 μm.

### 3.3 In vitro functional assays

To evaluate how GW release rates affected LXR activation, we focused on the LXR target gene Abca1, which has been well studied as a robust indicator of LXR-induced transcript expression [12]. In our study, we measured the effects of GW-NPs on macrophage *Abca1* mRNA transcript expression using quantitative reverse transcription PCR (RT-qPCR) and ABCA1 protein expression using Western blot. Given the possible heterogeneity and variability in macrophages from mouse to mouse, we isolated BMDM from C57BL/6J mice and performed independent LXR activation studies.

To evaluate the effects of controlled release, we treated macrophages with three different NPs containing the same amount of GW for 90 min to allow for cellular uptake. Afterward, cells were washed and replaced with fresh medium and incubated for up to 24 h, before measuring mRNA and protein expression levels of ABCA1. As controls, we also treated macrophages with free GW for 90 min, which was then either removed (similar to all NP treatments) or replenished for a continuous treatment. Independent of treatment, there was approximately the same level of *Abca1* transcript 2 h following the removal of the NPs and free GW (Figure 4A). However, by 6 h and peaking at 24 h, cells that were left exposed to free GW had higher amounts of Abca1 transcript (Figure 4B and C).

**Figure 4.**
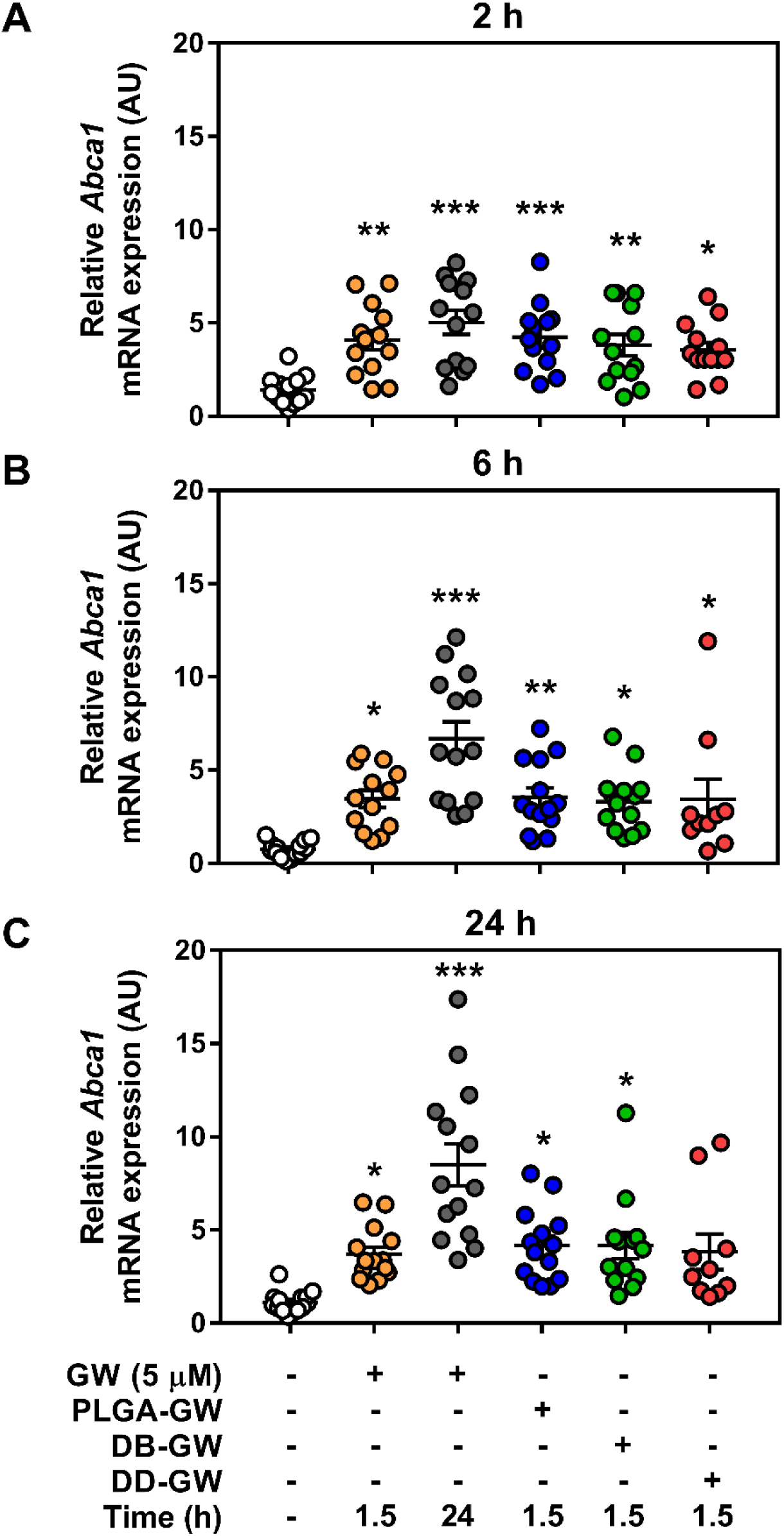
LXR target mRNA expression is unchanged by GW-NP addition. WT BMDM were treated with nanoparticles encapsulating 5 µM GW-3965 for 1.5 h. *Abca1* mRNA transcript expression was measured in samples A) 2 h, B) 6 h, or C) 24 h post-GW-NP treatment by RT-qPCR. Data represent mean ± SEM where *** p<0.001, ** p<0.01, and *p<0.05 are differences compared with vehicle control determined by one-way ANOVA (n = 10-13).

While all three NPs enhanced *Abca1* transcript expression compared to control, there were no differences between the NP-containing GW and the initial 90 min uptake of free GW at either 6 or 24 h. Despite this lack of change between NP formulations at the transcript level, ABCA1 protein amount was significantly higher in the DB-GW compared to the PLGA-GW and DD-GW conditions, which were both significantly higher than control but equal to free GW for 90 min (Figure 5).

**Figure 5.**
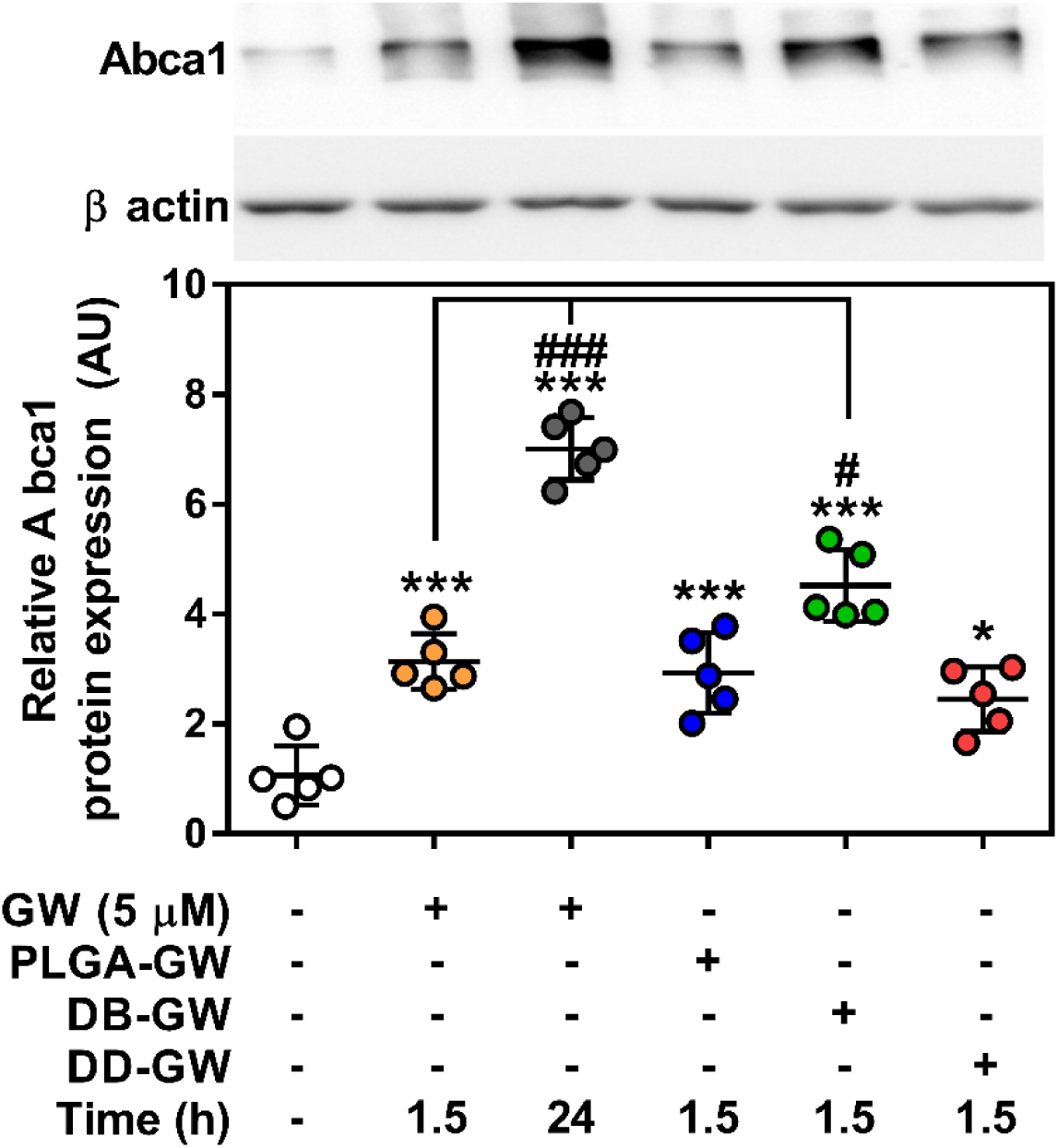
ABCA1 protein expression is upregulated with DB-GW nanoparticle treatment. WT BMDM were treated with nanoparticles encapsulating 5 µM GW-3965 for 1.5 h. ABCA1 protein was measured 24 h post-treatment and normalized to ß actin. Data represent mean ± SEM where *** p<0.001 and * p<0.05 are differences compared with vehicle control, while ### p<0.001 and # p<0.05 is compared to acute GW-3965 dose determined by one-way ANOVA (n = 5).

## 4. Discussion

Nanotechnology-based drug delivery platforms have revolutionized the landscape of drug development by increasing the blood circulation half-lives of drugs, decreasing their cytotoxicity, and delivering them to organ and tissues in a site-specific manner [16]. In addition, recent advances in biomaterials development led to endogenous and exogenous stimuli-responsive drug delivery platforms to enhance therapeutic efficacy [30]. In terms of endogenous stimuli, differences in pH and concentrations of reducing agents in the different microenvironments of the body are well studied and are currently being explored for therapeutic potential in clinical trials for cancer treatment [30]. However, stimuli-response drug delivery is less characterized in inflammation and cardiovascular disease applications. This may be due to the lack of available NP platforms with payloads that are well-characterized in cardiovascular and inflammation disease settings. The goal of this study was to explore if the differential release of drug through NP degradation due to endogenous stimuli would affect the drug target genes in macrophages, one of the primary cell types in atherosclerotic plaque. To this end, we developed and studied three different NP platforms containing LXR agonist GW-3965 and their effects in mouse macrophages.

Gadde et al, previously showed that PLGA-PEG nanoparticles containing GW attenuated the LPS induced inflammation in macrophages by downregulating ABCA1 expression [12]. In this study we used normal macrophages and all three GW-NPs upregulated the transcript expression of LXR target genes in macrophages and were in agreement with previous reports [12,14]. At the time points tested, there was no difference in *Abca1* mRNA expression between the formulations (Figure 4 and SI Figure 2). However, in the case of DB-GW NPs, ABCA1 protein expression was higher than the other NP treatments (Figure 5). Discrepancies in mRNA and protein expression of ABCA1 in macrophages has been observed previously [31], potentially due to extensive post-transcriptional regulation ABCA1 translation by a host of miRNAs [32]. While we did not investigate the possibility of differential miRNA expression in response to NP treatments, this could be a factor in the differences between mRNA and protein expression.

In addition, NP drug release can be influenced by a number of characteristics including polymer molecular weight, NP size and surface charge, cellular uptake pathways, intracellular components and, most importantly, cell types. In vivo and in vitro characterization of redox-responsive NPs have been primarily reported in tumor cells and tumor microenvironments, wherein cellular antioxidants, made up primarily of reduced glutathione (GSH), are found in much higher concentration than in extracellular fluids. However, we reasoned that GSH concentrations were not elevated in BMDM when compared to tumor cells, explaining the similar LXR target gene activation between slow releasing (PLGA) and redox responsive (DB and DD) NPs. However, in the context of atherosclerosis, oxidized LDL has been shown to increased intracellular GSH levels in macrophages in the atherosclerotic plaques [33]. It would be interesting in the future to test these conditions, as it may be possible that redox-reactive NPs would be beneficial for drug release. Also, there is the potential that the polymers themselves have elicited a ROS-generating response, diminishing cellular GSH:GSSG, and subsequently the rate of polymer degradation. Indeed, engineered NPs that use SiO_2_, Fe_3_O_4_, or CoO have been shown to stimulate ROS production in macrophages by diverse mechanisms [34].

Cholesterol regulation modulators, specifically reverse cholesterol transport enhancers have several therapeutic applications in atherosclerosis and heart disease settings. Recently, PLA-PEG NPs containing the anti-diabetic drug rosiglitazone (RSG) was shown to diminish inflammatory signaling in RAW264.7 macrophages [35]. Given the adverse whole-body effects of RSG administration [36], nanoparticle encapsulation presents an attractive targeting option to maximize tissue-specific benefits. Similar to RSG, whole-body pan-LXR activation has been shown to induce hepatic and adipose lipogenic gene expression, ultimately increasing plasma triglycerides and promoting steatosis of the liver [37]. Therefore, macrophage-specific nanoparticle targeting has been an area of active research in recent years.

## 5. Conclusions

Our results demonstrate that these two redox-responsive NP formulations were effectively taken up in vitro by macrophages and activated transcription of the LXR target gene, Abca1. Further research is warranted to assess NP response to greater cellular redox potential, as well as conditions mimicking cellular stress, in order to determine their effectiveness as therapeutic vehicles for diseases such as atherosclerosis.

## Supplementary Materials

The following are available online at www.mdpi.com/xxx/s1, Figure S1: Schematic diagram of DB and DD chemical structures and synthesis, Figure S2: Particle size and surface charge of empty NPs, Figure S3: Abca1 mRNA time course reveals no change in expression.

**SI Figure 1.**
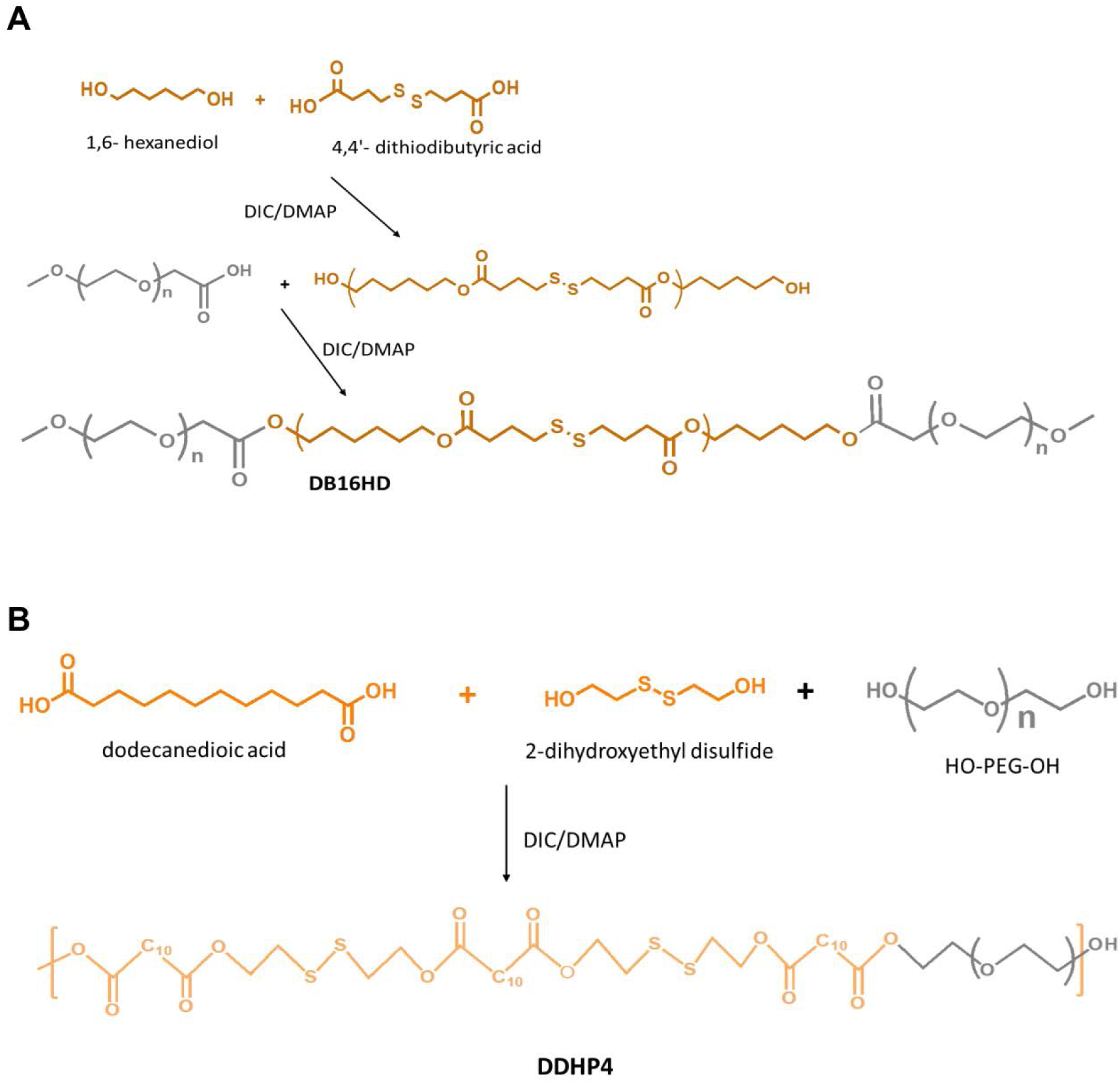
Schematic diagram of DB and DD chemical structures and synthesis.

**SI Figure 2.**
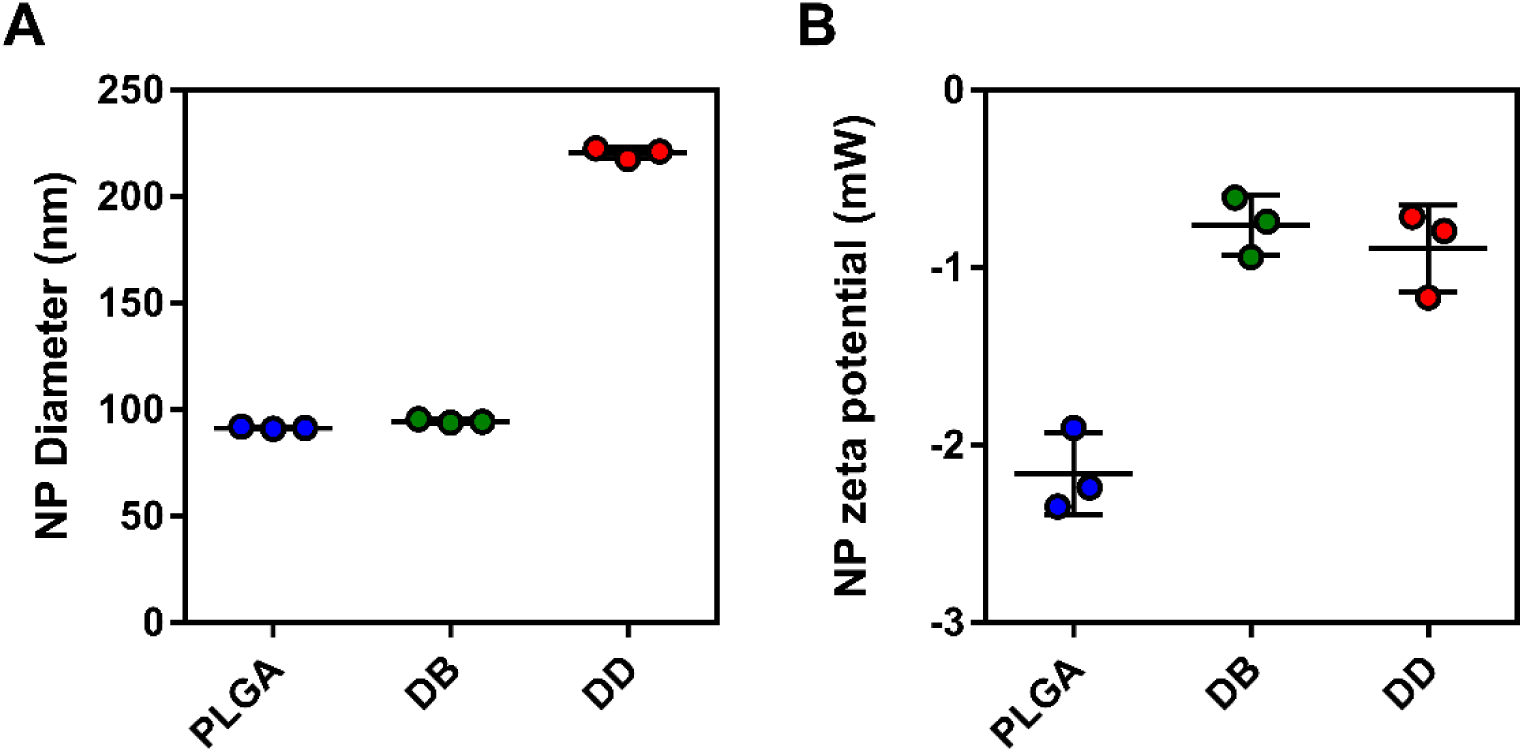
Empty NP characteristics. A) Particle size and B) surface charge. Each data point represents the mean of all values within each time point ± SEM (n = 3).

**SI Figure 3.**
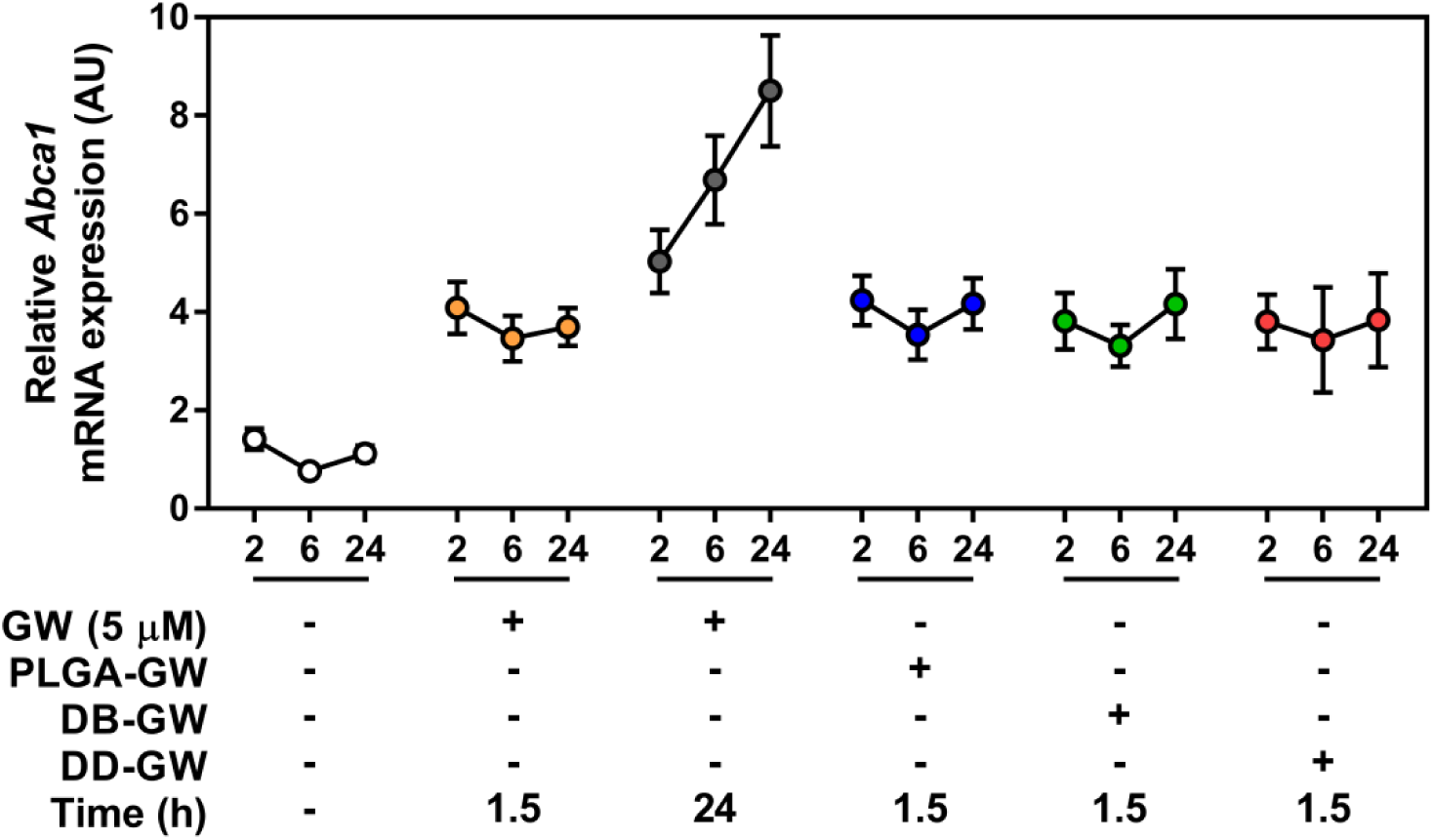
*Abca1* mRNA time course reveals no change in expression. Time course depicting data from Figure 4. Each data point represents the mean of all values within each time point ± SEM (n = 10-13).

## Author Contributions

Conceptualization, Tyler K.T. Smith, Suresh Gadde and Morgan D. Fullerton; Formal analysis, Tyler K.T. Smith, Zaina Kahiel, Suresh Gadde and Morgan D. Fullerton; Funding acquisition, Morgan D. Fullerton; Investigation, Tyler K.T. Smith, Zaina Kahiel, Nicholas D. LeBlond, Peyman Ghorbani, Eliya Farah and Suresh Gadde; Resources, Marceline Cote; Supervision, Suresh Gadde and Morgan D. Fullerton; Writing – original draft, Tyler K.T. Smith, Suresh Gadde and Morgan D. Fullerton; Writing – review & editing, Tyler K.T. Smith, Zaina Kahiel, Nicholas D. LeBlond, Peyman Ghorbani, Eliya Farah, Marceline Cote, Suresh Gadde and Morgan D. Fullerton.

## Acknowledgments

We would like to thank the lab of Dr. Jean-François Couture for reagents.

